# Genome-wide identification of notochord enhancers comprising the regulatory landscape of the *Brachyury* (*T*) locus in mouse

**DOI:** 10.1101/2023.06.21.545916

**Authors:** Dennis Schifferl, Manuela Scholze-Wittler, Alba Villaronga Luque, Milena Pustet, Lars Wittler, Jesse V. Veenvliet, Frederic Koch, Bernhard G. Herrmann

## Abstract

The node and notochord are important signaling centers organizing dorso-ventral patterning of cells arising from neuro-mesodermal progenitors forming the embryonic body anlage. Due to the scarcity of notochordal progenitors and notochord cells, a comprehensive identification of regulatory elements driving notochord-specific gene expression has been lacking. Here we have used ATAC-seq analysis of FACS-purified notochordal cells from TS12-13 mouse embryos to identify 8921 putative notochord enhancers. In addition, we established a new model for generating notochordal cells in culture, and found 3728 of these enhancers occupied by the essential notochordal regulators Brachyury (T) and/or Foxa2. We describe the regulatory landscape of the T locus comprising 10 putative enhancers occupied by these factors and confirmed the regulatory activity of 3 of these elements. Moreover, we characterized one new notochord enhancer, termed *TNE2*, in embryos. *TNE2* complements the loss of *TNE* in the trunk notochord, and is essential for notochordal cell proliferation and differentiation in the tail. Our data demonstrate the essential role of Foxa2 in switching T expressing cells from a NMP/mesodermal trajectory to the notochordal fate.

**Summary statement:** Combining multi-omics assays of purified embryonic and in vitro generated cells we identified thousands of notochord enhancers comprising *TNE2* essential for *T* expression and tail development of the mouse embryo.

## Introduction

The notochord is a rod-like structure situated below the neural tube and extending all along the trunk and tail of the embryo. As the defining characteristic of chordates, it serves two major functions: to provide structural stability to the developing embryo and signals for patterning of neighboring tissues (Stemple, 2005). It comprises an epithelium wrapped around a core of vacuolated cells, which makes the structure both solid and laterally flexible. In aquatic animals, these characteristics are essential and allow the embryo to swim. In fact, in some primitive fish the notochord serves as the main axial skeleton. In most chordates and all mammals, however, the notochord is a transient embryonic signaling center, which regresses during fetal development and forms the nucleus pulposus integrated in the intervertebral discs (Choi et al., 2008; McCann et al., 2012).

Adjoining organ anlagen of all three germ layers, the notochord lies ventral to the neuroectoderm, dorsal to the gut endoderm, and is flanked by somitic mesoderm. Patterning of the neural tube via secretion of Hedgehog ligands and Bmp antagonists from the notochord is a well-described signaling function (Briscoe and Ericson, 1999; McMahon et al., 1998). Further, the node induces the left-right axis and the notochord provides signals for patterning the endoderm during organogenesis (Cleaver and Krieg, 2001; Shiratori and Hamada, 2006). In the mesodermal lineage, the notochord is required for somite patterning, (Brand-Saberi et al., 1993; Fan and Tessier-Lavigne, 1994) sclerotome induction, vertebral column differentiation and segmentation (Ward et al., 2018).

Axial mesoderm emerges during gastrulation and is constituted of the prechordal plate, the anterior head process, and the node, which gives rise to trunk and tail notochord (Sulik et al., 1994; Tam and Beddington, 1987; Wymeersch et al., 2019; Yamanaka et al., 2007). Morphogenesis of these structures is controlled by a set of developmental transcription factors, in particular Foxa2 and Brachyury (T; Tbxt). The endodermal master regulator Foxa2 is essential for notochord specification at all axial levels (Ang and Rossant, 1994; Weinstein et al., 1994). Node formation and specification of trunk and tail notochord is controlled by the pan-mesodermal regulator Brachyury in a dosage dependent manner (Herrmann, 1995; Herrmann et al., 1990; Stott et al., 1993). Noto is required for tail notochord and disrupted in the truncate mutant, which displays a shortened tail phenotype (Abdelkhalek et al., 2004; Plouhinec et al., 2004; Zizic Mitrecic et al., 2010). During trunk notochord morphogenesis, Noto functions synergistically with Foxa2, before it becomes essential for tail notochord maintenance (Yamanaka et al., 2007). Of the three TFs, Noto is the only factor that is exclusively expressed in axial mesoderm precursors from E7.5. _3_

The combined activities of T, Foxa2, Noto and other transcription factors establish the gene regulatory network that govern notochord morphogenesis (Di Gregorio, 2020; Matsumoto et al., 2007; Passamaneck et al., 2009; Song et al., 2023; Yamanaka et al., 2007). So far, few notochord enhancers have been identified in mice, including *TNE*, *Foxa2 NE*, *NOCE* and *Sfpe2* at the *T*, *Foxa2*, *Noto* and *Shh* loci respectively (Alten et al., 2012; Jeong and Epstein, 2003; Nishizaki et al., 2001; Schifferl et al., 2021). We hypothesized that in addition to *TNE*, which is active during trunk notochord specification and essential for tail development, a second enhancer compensating for its loss must be located upstream of the *T* gene (Schifferl et al., 2021). Enhancers can be predicted by assessing chromatin accessibility, indicative histone modifications and transcription factor binding (Heintzman et al., 2007; Rada-Iglesias et al., 2011; Visel et al., 2009). Previous studies on notochord enhancers were limited due to the small number of cells available from embryonic material and the restricted accessibility of axial mesoderm (Tamplin et al., 2011).

In this study, we present a comprehensive and integrated approach utilizing ATAC-seq, ChIP-seq, and transcriptome profiling to identify notochord enhancers throughout the genome and their corresponding target genes. We elucidate the cis-regulatory landscape of the *T* locus comprising multiple enhancers, and identify crucial enhancers required for *T* expression in the notochord and essential for notochord formation and differentiation.

## Results and discussion

### Genome-wide identification of notochord-specific enhancers bound by T and/or Foxa2

The notochord is characterized by co-expression of Brachyury and Foxa2, which are also expressed in NMPs and mesoderm or endoderm, respectively. Noto, on the other hand, is notochord-specific. To characterize the enhancer landscape of the notochord we engineered a Noto::H2B-mCherry/T::Venus/Foxa2::mTurquoise triple reporter mESC line allowing the isolation of putative notochord (Noto+/T+/Foxa2+) progenitor cells (NotoP; Fig. 1A). The reporter line was used for the generation of embryos via diploid morula aggregation (Fig. 1B). Caudal ends of E8.5 and E9.5 embryos were dissected at the somite border, and Noto^mC+^/T^Ve+^/Foxa2^mT+^ cells were purified by flow cytometry (Fig. 1C). For comparison, we also purified trunk notochord (Fig. S1A) and paraxial mesoderm progenitors (MP; T^Ve+^/Noto^mC-^/Foxa2^mT-^; Fig.1C). Cell pools of Theiler stage 12 and 13 were used for ATAC-seq, E8.5 and E9.5 derived cells for transcriptome profiling (Fig.1C, Fig. S1A-D).

**Figure 1.**
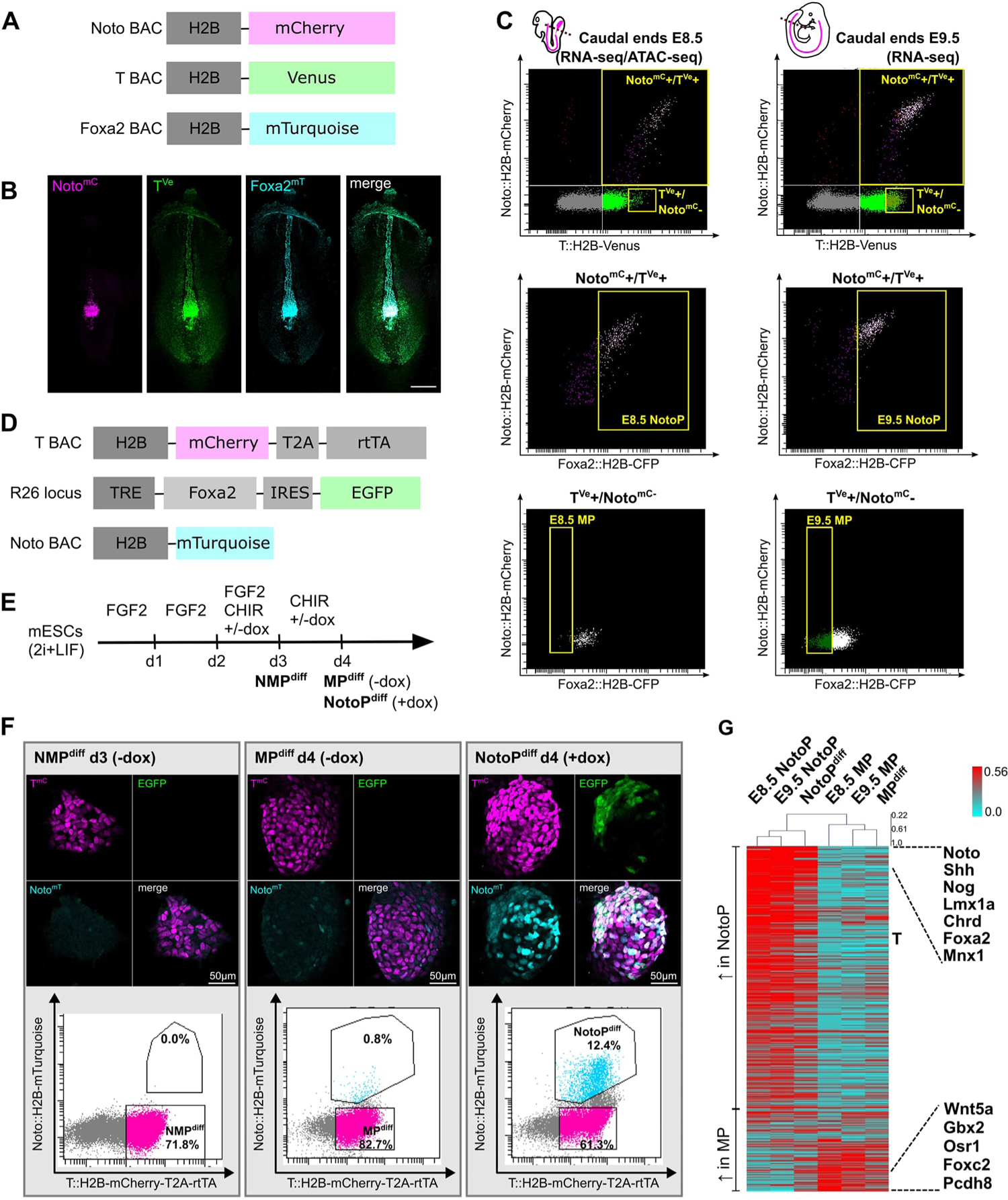
Derivation of NotoP and MP progenitor cells *in vivo* and *in vitro*. **(A)** BAC constructs integrated into a mESC line for the establishment of a Noto^mC^/T^Ve^/Foxa2^mT^ triple reporter line. **(B)** E8.5 Noto^mC^/T^Ve^/Foxa2^mT^ mouse embryo. Maximum intensity projection of images acquired by confocal microscopy. Scale bar = 200 µm. **(C)** FACS gating for purification of NotoPs and MPs from caudal ends of E8.5 and E9.5 embryos dissected at the somite border. **(D)** Genetic modifications of a mESC line used for T driven, DOX inducible overexpression of Foxa2. **(E)** Differentiation scheme for the *in vitro* generation of NMPs, MPs and NotoPs from mESCs carrying the modifications shown in (D); modified from Gouti et al 2017. **(F)** Images of differentiated colonies acquired by confocal microscopy, and corresponding FACS profiles. **(G)** Ranked heat map showing per gene normalized FPKM values of 418 differentially expressed genes (3-fold change in NotoPs vs. MPs at both E8.5 and E9.5) and corresponding *in vitro* generated cells (NotoP^diff^; MP^diff^). Genes sorted by average fold change in E8.5 and E9.5 samples. Examples of notochord (top) and mesoderm (bottom) progenitor marker genes are indicated on the right.

Embryonic notochords do not provide sufficient cell numbers for generating ChIP-seq data from FACS-sorted material. To overcome this limitation, we developed an *in vitro* model for differentiation of mESCs into NotoPs. In parallel we produced MPs under similar experimental conditions. Notochordal cells require high levels of T in combination with Foxa2. Therefore, we generated an mESC line allowing doxycycline-inducible expression of Foxa2 and eGFP, driven by reverse tetracycline controlled transactivator (rtTA; (Gossen et al., 1995) expressed under control of T. In addition, this line carries a Noto^mT^ reporter (Fig. 1D). We followed a well-established protocol for generating neuro-mesodermal progenitors (NMPs) on day 3 of culture (Fig. 1E; (Gouti et al., 2014). CHIR treatment for one more day generated MPs (herein termed MP^diff^ to distinguish them from embryonic MPs), as previously shown. However, parallel treatment with CHIR and Dox for 2 days (d2-d4) caused Foxa2 and eGFP expression, and efficiently generated Noto^mT+^ cells, as shown by FACS and fluorescent microscopy (Fig. 1F; Fig. S2A). To test if these cells resemble embryonic NotoPs, we isolated Noto^mT+^/T^mC+^ cells (Fig. 1F) and performed RNA-seq. In parallel we analyzed MP^diff^ cells (T^mC+^/Noto^mT-^). We compared the transcriptomes of *in vitro* generated to embryonic cells isolated from E8.5 and E9.5 embryos. For embryonic NotoP and MP cells, we identified 319 and 109 specifically expressed genes, respectively (3-fold change of FPKM value in NotoP vs. MP at E8.5 and E9.5) comprising many known markers for both cell types (Fig. 1G, Fig. S2B, Table S1). The comparison revealed striking similarity between the expression profiles of embryonic and *in vitro* generated cells. Immunofluorescent staining and qPCR confirmed that T and Foxa2 are upregulated in Noto^mT+^/T^mC+^ cells, while expression of the paraxial mesoderm marker *Tbx6* is low (Fig. S2C-D). We conclude that our *in vitro* model is suitable for efficiently generating notochord-like cells strongly resembling embryonic NotoPs.

Next, we identified putative notochord enhancers. We performed ATAC-seq on embryonic NotoPs and MPs. Differential peak detection identified 8921 open regions outside of promoters (+/- 5 kb from TSS) with higher accessibility in NotoP and 4876 regions in MP cells (Fig. 2A; Fig. S3A). To characterize these accessible regions with respect to T and/or Foxa2 binding we generated cultures enriched for NotoP^diff^ cells (CHIR+; Dox+; d4). We performed ChIP-seq for T and Foxa2 on bulk cultures. In addition, for determining specific chromatin signatures we FACS-purified NotoP^diff^ (CHIR+; Dox+; d4) and MP^diff^ (CHIR+; Dox-; d4) cells as above and analyzed several histone marks (H3K27ac, H3K4me3, H3K4me1 and H3K9me2) via ChIP-seq. For comparison we used T ChIP-seq data previously generated from *in vitro* differentiated NMPs (Koch et al., 2017).

**Figure 2.**
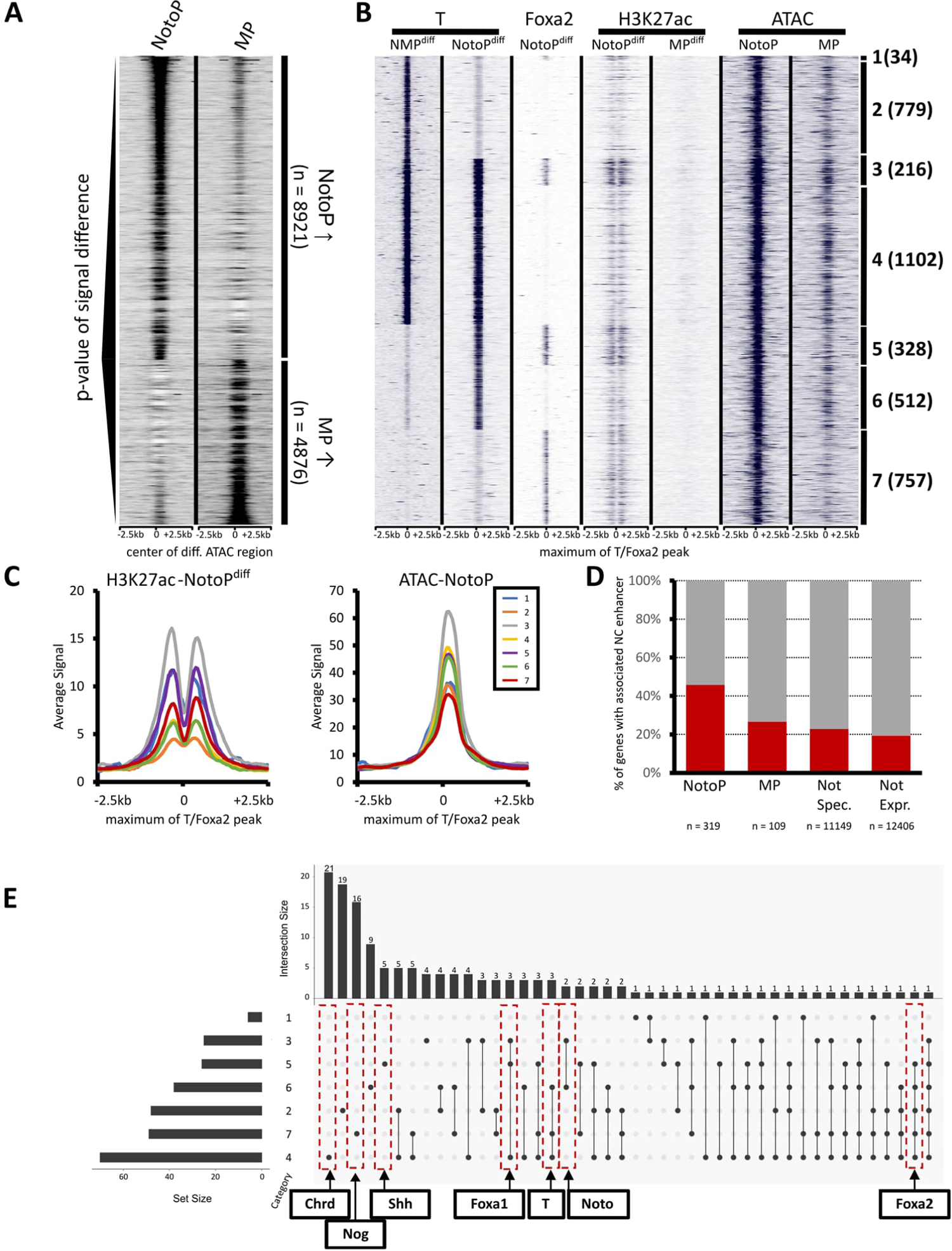
Genome-wide prediction and analysis of notochord enhancers. **(A)** Heatmap showing ATAC-seq signals of NotoP and MP cells around 8921 notochord enhancer candidates defined by differential peak detection, sorted by p-values. **(B)** Heatmap of 3728 differential NotoP accessible sites bound by T and/or Foxa2 in NotoP^diff^ cells. Putative enhancers are categorized into all possible combinations of T and Foxa2 binding combinations (1-7, right) and the number of putative enhancers per category is shown in parentheses. **(C)** Average profiles of H3K27ac and ATAC-seq signal for all enhancer categories. **(D)** Barplot quantifying the association of putative notochord enhancers to NotoP, MP, not specifically expressed and not expressed genes. **(E)** UpSet plot for the 146 NotoP specific genes showing their associated enhancers according to categories 1-7.

The co-expression of T and Foxa2 in our *in vitro* model strikingly changed the chromatin landscape from a mesoderm signature (CHIR+; Dox-; d4) to a notochord signature (CHIR+; Dox+; d4) (Fig. 2B, Fig. S3B). Among the 8921 open regions differentially accessible in NotoPs versus MPs we identified 3728 sites bound by T and/or Foxa2 in NotoP^diff^ cells (Fig. 2B; Fig. S3B). Using H3K27ac data as strong indicator of enhancer activity (Creyghton et al., 2010) we found the highest signal in enhancers binding both T and Foxa2 (category (Cat) 3, followed by 5 and 1; Fig. 2C). Cat 3 enhancers also show the highest ATAC signal in embryonic NotoPs, and are as well bound by T in NMPs. Cat 5 enhancers specifically occupied by T and Foxa2 in NotoP^diff^ cells show lower ATAC and H3K27ac signal. This suggests that Foxa2 and T could have additive effects on enhancer activation. On the other hand, based on the H3K27ac signal, Foxa2 binding alone (Cat 7) seems to exert higher enhancer activity than T alone (Cat 2, 4, 6), whereas the accessibility of cat. 4, 5 and 6 enhancers appears to be equal. A considerable fraction (2393/3728; 64.2%) of notochord enhancers showed T binding alone (Cat 2, 4 and 6), a smaller fraction (757/3728; 20.3%) only Foxa2 binding (Cat 7). Most of the T binding enhancers (2131/3728; 57.1%) are also T bound in NMPs, while 840 (22.5%) T bound enhancers were only occupied in notochord cells. Thus, Foxa2 co-expression with T opens enhancers not occupied by T when Foxa2 is not expressed, and thus this cooperative action changes the genomic landscape to a notochord signature.

Next, we determined the enhancer presence in the vicinity of 319 genes specifically upregulated in NotoP cells (Fig. 1G). We found at least 1 notochord enhancer each in the gene bodies or genomic intervals comprising the intergenic regions and gene bodies of the immediate neighbors of 146 (46.4%) of these genes (Fig. 2D). Figure 2E lists the enhancers according to cat 1-7 found next to the 146 genes, showing examples of important well-known notochord marker genes. Strikingly, most of these genes are associated to several putative enhancers falling into different enhancer categories and none of them are associated to a T-NMP^diff^ peak (Cat 2) alone. Genome browser screen shots of important markers are shown on Figure S4, housekeeping genes for control on Figure S5. For instance, in addition to notochord enhancers described previously, we found new enhancers for *Noto, Foxa2, Chrd, Nog and Foxa1* (Schifferl et al 2021, Alten et al 2012, Nishizaki et al 2005, Jeong and Epstein, 2003). A listing of all notochord enhancers identified here and their associated genes as well as the list of the 319 NotoP genes and the categories of their associated enhancers are listed in Table S2 and Table S3 respectively. Our data provide an important resource of novel putative notochord enhancers.

### The regulatory landscape of the *T* locus in notochord and mesoderm

Next, we used our integrated ATAC and ChIP data to further characterize the regulatory landscape of the *T* locus in notochord and mesoderm. We previously identified the *TNE* enhancer, which is critical for tail notochord development, but does not fully explain the severe loss-of notochord and tailbud phenotype of a 37 kb deletion (T^UD^), suggesting additional notochord and mesoderm elements are present in this region (Schifferl et al., 2021). Our ATAC-seq data of the *T* locus shows 10 peaks including *TNE* (Fig. 3).

**Figure 3.**
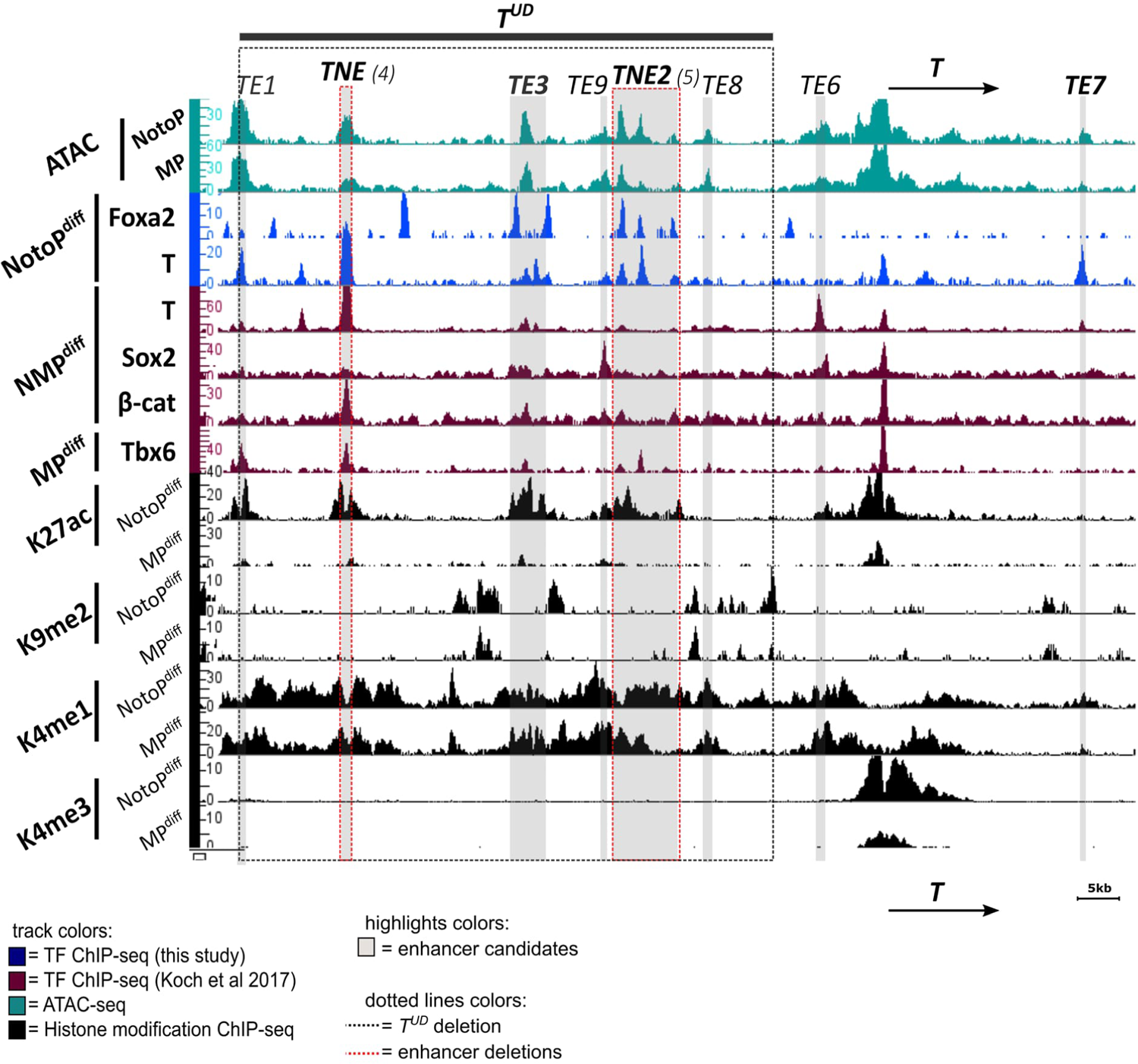
Identification of putative enhancers at the mouse *T* locus. Genome browser snapshot showing the mouse *T* locus with signal density tracks of ATAC-seq, TF ChIP-seq, and Histone 3 ChIP-seq data. Track maxima are normalized to the number of mapped reads for each antibody. Enhancers are highlighted in grey. The *T^UD^* and *TNE* regions were described previously (Schifferl et al 2021). The number in brackets indicates the enhancer category as shown on Fig.2B.

Three ATAC-seq peaks are bound by T and Foxa2 and carry the active enhancer mark H3K27ac in NotoP^diff^ cells (Fig. 3). Therefore, we used a 4.4 kb fragment encompassing these T and Foxa2 binding sites to perform an enhancer assay. The embryonic activity assay at E9.0 shows Venus reporter expression in the posterior notochord, decreasing towards the head, identifying the 4.4 kb fragment as a notochord-specific enhancer (Fig S6A). We designated this enhancer *T locus notochord enhancer* 2 (*TNE2*).

In addition, based on ATAC peaks and H3K27 acetylation data we define several more candidate *T* locus enhancers, *TE1*, *TE3*, *TE9* and *TE8* within the *T^UD^* region (Schifferl et al. 2021), as well as *TE6* −5kb upstream and *TE*7 13 kb downstream of the *T* transcription start site (Fig.3). We integrated ChIP-seq data for T, Sox2, beta-Catenin and Tbx6 published previously, to relate these enhancers to regulatory processes occuring in NMPs and during mesoderm formation (Koch et al. 2017). For example, Sox2 acting antagonistically to T in the neural versus mesodermal lineage choice, binds to *TE6* and *TE9* in NMPs, while the WNT signal mediator beta-catenin, which cooperates with T in NMPs and NotoPs, binds to *TE3* and *TNE*. In contrast, Tbx6, which has a repressive effect on NMP maintenance and *T* expression, is detected at *TE1, TNE, TNE2* and possibly *TE3*. All four TFs also bind at the *T* promoter.

We assayed the activity of *TE3*, characterized by T peaks flanked by Foxa2 binding in NotoP^diff^ cells using the HSP68-Venus reporter. This element showed expression in the tailbud mesoderm, posterior neural tube and gut of the tail at E10.5, but not in the notochord (Fig S6B). This data suggests that *TE3* acts as enhancer in tailbud NMPs. The significance of the flanking Foxa2 peaks is unclear.

Moreover, we assayed the putative enhancer *TE7*, which is located downstream of the *T* gene and was not identified by chromatin marks or differential ATAC analysis, but showed a small ATAC peak and T binding in NotoP^diff^ cells. Beta-Gal reporter activity was detected in the entire notochord of E9.0 embryos, albeit with increasing staining towards the head (Fig S6C). Thus, *TE7* identifies another notochord enhancer. The data suggest that the activity of *TE7* is increasing during notochord differentiation, which might explain that it was not detected in our differential ATAC-seq data derived from caudal end NotoPs.

### A new *T* locus notochord enhancer, *TNE2*, complements and synergizes with *TNE* in notochord development

Next, we investigated the function of several putative *T* locus enhancers by employing the CIRSPR/Cas9 system to generate a series of knockout mESC lines and embryos carrying the *Noto::H2B-mCherry* (Noto^mC^) reporter (Fig. S7, Table S4). We generated mutant embryos by tetraploid complementation assays and evaluated the phenotype between E9.5 and E12.5 by visualizing Noto^mC^ reporter and T as well as Sox2 protein expression. Deletions of *TE1*, *TE3*, *TE9 and TE*7 did not result in embryonic defects enhancing the phenotypes of the corresponding parental line, which contained either the wild type *T* locus or the heterozygous *T^LD^*deletion covering the entire *T* locus (Schifferl et al., 2021). The data suggest that these cis control elements either have no critical function or are redundant (Table S5).

However, the homozygous deletion of *TNE2* resulted in a notochord deficiency phenotype. In E9.75 *T^ΔTNE2/ΔTNE2^* mutant embryos, less T protein is detected in the trunk notochord marked by Noto^mC^ than in wild type embryos, and the notochord is partially disrupted at the hindlimb level (Fig. 4 A,C). At E11.5, the Noto^mC^+ notochord progenitor domain is present in the outgrowing tailbud, but Noto^mC^+ cells in the midline are dwindling away quickly towards the anterior and the notochord is not formed (Fig. 4 B,D). The reduced T protein level in the trunk notochord apparently is still sufficient for trunk development, while tail notochord formation is not supported resulting in a tailless phenotype at E12.5 (Fig. 4 G-H’’’).

**Figure 4.**
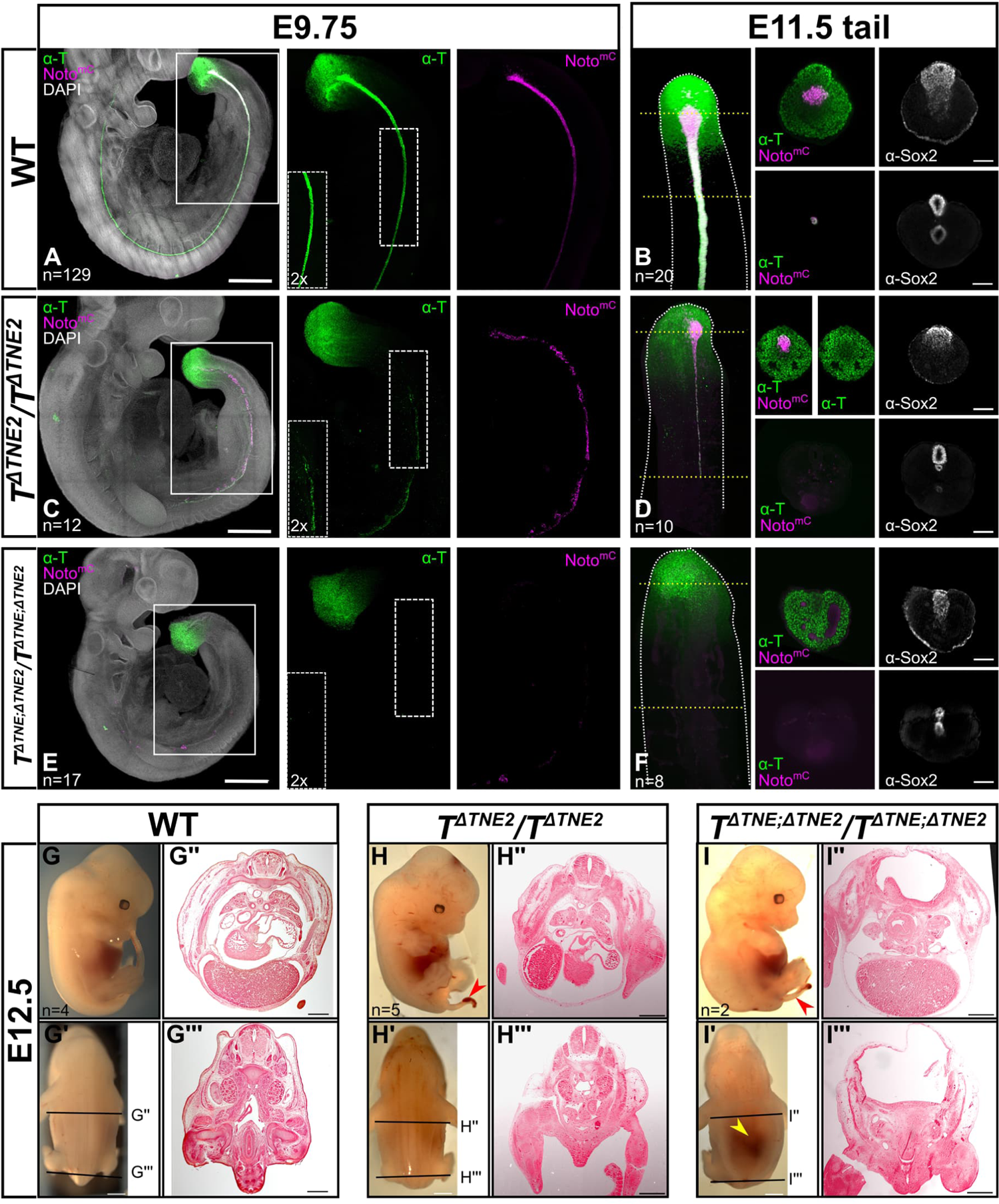
*TNE2* is an essential notochord enhancer of *Brachyury* and interacts with *TNE*. **(A,C,E)** Maximum intensity projections of E9.75 embryos with Noto^mC^ reporter signal, immunostaining for T (green) and DAPI nuclear staining (grey). Scale bar = 500 µm. The square indicates the area magnified in single channels. 2x enhanced signal is shown in the bottom left panel of the T channel. **(B,D,F)** Maximum intensity projections of E11.5 tails with immunostaining for T (green) and the Noto^mC^ reporter signal (magenta). Yellow dashed lines indicate the position of optical sections: Scale bar = 100 µm. **(G-I)** Lateral and dorsal views of E12.5 embryos with histological sections at the indicated axial levels. Scale bar (lateral and dorsal views) = 1 mm. Scale bar (histological sections) = 500 µm. Red and yellow arrows indicate the tail phenotype and neural tube defects respectively.

In E9.75 *T^ΔTNE; ΔTNE2/ΔTNE; ΔTNE2^* double knockout embryos, T expression was detected in the tailbud, but not in notochord progenitors, and the notochord was not formed (Fig. 4E). Few cells with weak Noto^mC^+ signal were visible in gut endoderm, indicating improper specification (Fig. 4E). No Noto^mC^+ cells nor a notochord were detected in the tailbud at E11.5 (Fig. 4F). Immunofluorescent staining for Olig2 and Nkx2.2 revealed neural tube patterning defects (Fig. S8). Nkx2.2 expression was shifted ventrally, while Olig2 was not detected. Consistent with the latter, neural tube closure and differentiation was severely affected at E12.5, resulting in embryonic lethality (Fig. 4I-I’’’).

Our data show that *TNE* can compensate for the loss of *TNE2* during trunk notochord formation and vice versa (Schifferl et al., 2021). The double KO phenotype shows that the combined activity of these two enhancers is required and sufficient for notochord formation and differentiation in the trunk and tail, at least until the end of axial elongation. *TE7* is not able to compensate for the loss of *TNE* and *TNE2*. The reduced T protein level in either KO suggests that the two enhancers act additively during trunk notochord development. The partial disruption of the notochord in the posterior trunk of either single KO suggest that this region already requires a higher level of T expression for supporting normal notochord development. Tail notochord formation, however, does not take place at all without both enhancer activities. The increasing requirement of T for notochord development along the axis has been reported previously (Stott et al. 1993, Herrmann 1995). The single loss of function phenotypes reveal different roles of *TNE* and *TNE2*, at least in the tail. There, *TNE* is mainly required for notochord progenitor maintenance, and possibly for notochordal cell proliferation and differentiation, since embryos lacking *TNE2* still form Noto^mC^+ notochord progenitor cells, while they are quickly fading out in the midline toward the anterior. *TNE2*, on the other hand, is required for proper notochord cell specification and differentiation, since embryos lacking *TNE* show a dispersion of Noto^mC^+ cells into all cell lineages of the tailbud, while the notochord is missing and the NotoP domain is shrinking during tail outgrowth (Schifferl et al., 2021). Of note, our data suggest cooperative binding of T and Foxa2 at *TNE2*, while *TNE* appears to bind T and beta-Catenin, consistent with the role of canonical Wnt-signaling in notochord specification (Ukita et al., 2009). Thus, T seems to act earlier in the lineage through *TNE*, followed by cooperative action with Foxa2 through *TNE2*. Our enhancer reporter assays suggest that *TE7* is active even later, upon further differentiation and possibly in the maintenance of the notochord, as during axis elongation the reporter activity is low in the posterior end and increasing towards the head.

The strong *T^UD^*homozygous deletion phenotype reported previously is, at least with respect to the lack of the notochord, explained by the loss of *TNE* and *TNE2*, while *TE7* is not involved (Schifferl et al., 2021). However, the missing tailbud in the deletion mutant is not explained, since *TNE/TNE2* double KO embryos still form the tailbud and show tail outgrowth. *TE3* driving expression in tailbud NMPs might be responsible for tail outgrowth in the absence of *TNE/TNE2*. However, our *TE3* deletion analysis did not show the expected loss of the tailbud in the absence of the enhancer, even in combination with the deletion of *TE9*, an enhancer candidate showing Sox2 and beta-Catenin binding in NMPs (Fig. S7; Table S5). Therefore, the control elements required for tailbud formation and tail outgrowth must remain undefined at this point.

Note: while this work was prepared for publication, complementary work dissecting the regulatory landscape of the *T* locus with respect to notochord control elements was reported on the bioRxiv preprint website (Kemmler et al., 2023). In this work, the notochord enhancers *TNE*, *TNE2* and *TE7* were termed *T3*, *C* and *I*, respectively. The genomic fragments defining these enhancers are different from ours though with considerable overlaps.

## Acknowledgements

We are grateful to the MPIMG transgenic unit and animal facility, especially Karol Macura, Judith Fiedler, Adrian Landsberger, Mirjam Peetz, Dijana Micić and Christin Franke for performing the morula aggregation experiments and animal husbandry. We thank Sandra Währisch and Gaby Bläß for tissue culture support, Claudia Giesecke-Thiel, Erwin Weiß and Uta Marchfelder from the MPIMG flow cytometry unit for FACS support and all members of the MPIMG microscopy and sequencing core facilities for technical support.

## Materials and methods

### General mESC procedures

All cell lines used in this study were derived from male mESCs of the G4 hybrid line 129S6/SvEvTac x C57BL/6Ncr (George et al., 2007). mESC clones were regularly tested for possible mycoplasma contamination, using PCR Mycoplasma Test Kit II (*Applichem A8994*) according to the manufacturer’s recommendations.

### Generation and integration of BAC transgenes

For the generation of reporter and driver transgenes, BACs containing ∼200 kb C57/BL6 genome surrounding the mouse *brachyury* (RP24-530D23), *Foxa2* (RP23-254G2) and *Noto* (RP23-289M19) genes were obtained from BACPAC resources. The *T::H2B-Venus*, *T::H2B-mCherry-T2A-irTA*, *Foxa2::H2B-mTurquoise* and *Noto::H2B-mTurquoise* BACS carrying neomycin, hygromycin, puromycin and blasticidin resistance cassettes, respectively, were engineered via Red/ET recombineering (Muyrers et al., 1999) as described previously (Schifferl et al., 2021). For random integration of BACs, 5µg of each construct were linearized using PI-SceI (*New England Biolabs R0696S*) and electroporated into 3×10^6^ mESCs. Approximately 30h after electroporation, selection was initiated as shown in Table S4. Selection medium was refreshed daily until single colonies were clearly visible after approximately one week. Single clones were picked and genotyped by PCR. Oligonucleotides used in this study are listed in Table S6.

### Generation of enhancer mutants

Homozygous *T^ΔNE2^/T^ΔNE2^* single and *T^ΔTNE; ΔNE2^/ T^ΔTNE; ΔNE2^* double mutants were generated from cells carrying the *Noto::H2B-mCherry* construct using CRISPR/Cas9 as reported on previously (Schifferl et al 2021).

### Recombinase mediated cassette exchange

For the generation of enhancer reporter and Foxa2 overexpression cell lines, 3×10^5^ mESCs with a modified Rosa26 harboring locus (Vidigal et al., 2010) were co-transfected with 5 µg of linearized NE2-HSP68-Venus, TE3-HSP68-Venus, TE7-HSP68-bGal or TRE-Foxa2-IRES-EGFP constructs and 1 µg PGK-iCre vector using Lipofectamine 2000 (*Invitrogen 11668027*). Cells were cultured in ES+LIF containing 350 µg/ml geneticin (*Thermo Fisher 10131027*) for selection for successful recombination resulting in a switch from hygromycin to neomycin resistance.

### Generation of transgenic embryos

Transgenic mouse embryos were generated by diploid or tetraploid morula aggregation by the transgenic unit of the Max Planck Institute for Molecular Genetics in Berlin as described previously (Eakin and Hadjantonakis, 2006). All animal experiments were performed according to local animal welfare laws and approved by local authorities (covered by LaGeSo licenses G0243/18 and G0247/13).

### Embryo isolation

Timed pregnant foster mice were euthanized by carbon dioxide application and cervical dislocation. For whole mount immunofluorescence and tissue clearing embryos were isolated from uteri in 4°C pre-cooled PBS, fixed using 4% PFA / PBS (*Sigma Aldrich P6148*) in 4 ml glass vials (*Wheaton 224882*) and processed as described previously (Schifferl et al 2021). For RNA-seq and ATAC-seq, embryos were isolated in M2 medium (*Sigma Aldrich MR-015P*). Samples were further dissected into the sub-regions of interest using forceps (*Dumont 11251-10*). Tissue samples were kept on ice in M2 medium and processed subsequently.

### Whole mount immunofluorescence and tissue clearing

Immunofluorescent staining and clearing procedures of E9.75-E11.5 embryos were performed as described previously (Schifferl et al. 2021). Used Antibodies are listed in Table S7.

### Whole mount β-Galactosidase staining

Embryos carrying the *TE7-HSP68-bGal* reporter were fixed for 30 min at 4°C and subsequently washed 3x for 15 minutes in Rinse Buffer (50mM EGTA, 0.1% deoxycholate, 0.2% NP-40, 20mM MgCl2 in DPBS) at room temperature. After rinsing, embryos were incubated in staining solution (50mM K3Fe (CN)6, 50 mM K4Fe(CN)6, 50 mM EGTA, 0.1% deoxycholate (100x), 0.2% NP-40 (100x), 0,2M MgCl2, 1 mg/mL X-gal in DPBS) at 37°C overnight. Stained embryos were washed 3x with PBS and stored in 4%PFA/PBS at 4°C for secondary fixation.

### Fluorescence activated cell sorting

For FACS of embryonic material, single cell suspensions were prepared adding 100µl Trypsin/EDTA to the sample. After incubation at room temperature for 5 min, trypsin was quenched by adding 200µl PBS 5% BSA (*Sigma Aldrich A8412*).

For FACS of cell cultures, cells were washed 2x PBS and dissociated by trypsinization at 37°C for 10min. Trypsin/EDTA was quenched using a double volume of 5%BSA/PBS, resuspended and kept on ice until further procedure.

All samples were immediately filtered (35µm mesh) and sorted on a FACS Aria II (*Becton Dickinson*) flow cytometer. For Transcriptome analysis, cells were sorted into 350µl RLT Plus buffer (*Qiagen 1053393*) containing 1% β-Mercaptoethanol (*Sigma Aldrich M6250*) in 1.5 ml low binding tubes (*Thermo Fisher 90410*) and stored at −80°C until further procedure. For ChIP, cells were sorted into PBS 5% BSA in BSA coated glass tubes.

### Histology

PFA fixed E12.5 embryos were dehydrated through an ethanol series in 30%, 50%, 2x 70% EtOH for 15 minutes each, processed in a MICROM STP 120 processor (*Microm 813150*) and embedded in paraffin (*Leica 3801320*) utilizing an EC 350-1 embedding station (*Microm*). Sections of 10 µm thickness were prepared using a rotary microtome (*Microm*, HM355S), transferred onto adhesion microscope slides (*Menzel K5800AMNZ72*) and dried o.n. at 37°C. Eosin (*Merck 109844*) counterstaining was performed according to standard procedures and specimens were mounted in Enthellan (*Sigma-Aldrich 107960*). Sections were imaged using an AxioZoom V16 stereomicroscope (*Zeiss*).

### Microscopy

Embryos were imaged using a Zeiss LSM880 laser scanning microscope with Airyscan detector or Zeiss Light sheet LS Z1 with appropriate filters for DAPI, mTurquoise, EGFP, Venus, mCherry, Alexa488 or Alexa647. For Light sheet microscopy, specimens were cleared and embedded in 1.5% low melting agarose (Sigma-Aldrich A9414)/PBS. Agarose columns containing the samples were cleared in RIMS overnight before acquisistion. Processing was performed using ZEN Blue/Black (*Zeiss*) software.

### *In vitro* differentiation

NMPs, MPs and NotoPs were derived from *T^mC-2A-irTA^/TRE::Foxa2/Noto^mT^*mESCs following an established protocol (Gouti et al., 2014) with previously described modifications (Koch et al., 2017). For NotoP generation, 1ng/ml doxycycline was applied from d2-d4.

### RNA-Seq

For transcriptome analysis of FACS purified subpopulations, total RNA was isolated from 250 cells (or less; see Table S8) using the RNeasy MinElute kit (*Qiagen 74204*). RNA extraction was performed according to the manufacturer’s protocol with an additional DNase digest step between two washes with 350µl RW1. Therefore, reaction mixes of 10 µl DNase I (*Qiagen 79254*) and 1µl (=10U) DNase I (*Roche 4716728001*) in 70µl buffer RDD (*Qiagen 1011132*) to were applied to the spin columns for 15 min incubation at room temperature. Membranes were air dried for 10 min to remove ethanol remains and successively eluted in 2x 20 µl EB.

Sequencing libraries were prepared using the Ovation SoLo RNA-seq system (*NuGen*) according to manufacturer’s recommendations, starting at step A.9 with 12 µl purified DNA. After each amplification step, libraries were quantified with the Qubit the High Sensibility DNA assay (*Thermo Fisher 12102*). Library size was validated using DNA High Sensitivity Bioanalyzer chips (*Agilent 5067-4626*).

cDNA library pools (150nmol in 15µl) with 16bp barcode length (8bp barcode + 8bp UMI) were sequenced on using the Illumina HiSeq4000 (for E8.5/E9.5 NotoP, MP and Noto trunk) or NextSeq 2000 (for Noto^diff^, MP^diff^, NMP^diff^) platforms.

Prior to mapping, the first 5 nucleotides of the forward and reverse reads were trimmed using fastx_trimmer (http://hannonlab.cshl.edu/fastx_toolkit/index.html) according to manufacturer’s instructions (NuGEN). The resulting reads were mapped to chromosomes 1-19, X, Y and M of the mouse mm10 genome using TopHat2 (v2.1.1) and bowtie (v1.2.2) (Kim et al., 2013; Langmead et al., 2009) and the RefSeq annotation in gtf format (UCSC), providing the options ‘--no-coverage-search --no-mixed --no-discordant -g1 --mate-inner-dist 250 --mate-std-dev 100 --library-type fr-secondstrand’. Read duplications resulting from the PCR amplification of the library were removed using the NuDup deduplication script provided by NuGEN (http://nugentechnologies.github.io/nudup/). Wiggle files were generated with BEDTools version 2.23.0 (Quinlan and Hall, 2010), converted to bigwig format and visualized in the Integrated Genome Browser (Freese et al., 2016). FPKM values were calculated using Cufflinks version 2.2.1 (Trapnell et al., 2013) with options -u--no-effective-length-correction -b. Using FPKM values, per gene normalization, generation of heatmaps and non-hierarchical clustering of samples was performed in MeV (Howe et al., 2011).

### ATAC-Seq

ATAC-seq experiments were conducted following the established protocol (Buenrostro et al., 2013). A total of 2000 cells were utilized per ATAC-seq experiment. After FACS sorting, the cells were centrifuged and the resulting pellets were resuspended in 50μl of lysis buffer (10mM Tris pH 7.4, 10mM NaCl, 3mM MgCl2, 0.1% NP-40) before being subjected to centrifugation at 500g and 4°C for 10 minutes. The supernatant was discarded, and each pellet was then resuspended in the transposition reaction mix (25μl 2x TD buffer, 2.5μl Tn5 transposase, 22μl H2O), followed by incubation at 37°C for 30 minutes. Subsequently, the reaction was halted by adding PB buffer (*Qiagen*), and the tagmented DNA was purified using the MinElute kit (*Qiagen*). The purified DNA was combined with ATAC index PCR primers and 2x Kapa HiFi Hotstart Readymix and subjected to pre-amplification (98°C for 30 seconds, followed by 8 cycles of 98°C for 10 seconds, 63°C for 30 seconds, and 72°C for 1 minute) in a 50μl reaction volume. To determine the optimal number of additional cycles and prevent overamplification, 5μl of the preamplification mix was combined with primers, 1x Evagreen Sybr green (*Jena Biosciences*), and 2x Kapa HiFi Hotstart Readymix in a 15μl total volume and subjected to 30 cycles on a StepOne Plus instrument. The remaining pre-amplified samples (45μl) were subjected to an additional 7 (NotoP sample) or 6 (MP sample) cycles, resulting in a total of 15 (NotoP sample) or 14 (MP sample) cycles. The libraries were purified using MiElute columns (*Qiagen*), and the concentration was determined using the DNA HS Qubit assay (*Life Technologies*). Approximately 4ng of each library was subjected to DNA HS Bioanalyzer chip (*Agilent*) to assess library size and calculate molarities. The samples were pooled and subjected to paired-end sequencing on an Illumina NovaSeq 6000 platform with 2×100 bp read length.

### ChIP-Seq

For histone ChIP-Seq on sorted NotoP^diff^ and MP^diff^ cells, the iDeal ChIP-Seq kit (Diagenode) was used according to manufacturer’s instructions. Approximately 200,000 cells were used per ChIP. ChIP on bulk d4 *in vitro* differentiated notochord-like cells for the identification of T and Foxa2 binding sites was performed as described previously (Koch et al., 2011). Used Antibodies are listed in Table S7. ChIP-Seq sequencing libraries were generated using the TrueSeq ChIP-Seq kit (*Ilumina*) following the manufacturer’s instructions with minor modifications. After adapter ligation, 0.95x of AMPure XP beads (*Beckman Coulter A63880*) were used for a single purification and the DNA was eluted using 15 µl of resuspension buffer (RSB, Illumina). After the addition of 1 µl primer mix (25 mM each, Primer 1: 5’-AATGATACGGCGACCACCGA*G-3’; Primer2: 5’-CAAGCAGAAGACGGCATACGA*G-3’) and 15µl 2x Kapa HiFi HotStart Ready Mix (Kapa Biosystems), amplification was performed for 45 s at 98°C, 5 cycles of [15 seconds at 98°C, 30 s at 63°C and 30 s at 72°C] and a final 1 min incubation at 72°C. The PCR products were purified using 0.95x of beads and eluted using 21 µl of RSB. Libraries were directly amplified for an additional 13 cycles and purified using AMPure XP beads. The libraries were quantified using the Qubit DNA HS assay and the library size was validated using DNA HS bioanalyzer chips (*Agilent 5067-4626*). The samples were pooled and subjected to paired-end sequencing on an Illumina NextSeq 500 platform with 2×75 bp read length.

Reads were mapped to chromosomes 1-19, X, Y and M of the mouse mm10 genome using bowtie version 1.3.1 (Langmead et al., 2009), providing the options ‘ -y -m 1 -S -I 100 -X 500’. The mapping information of the paired-end reads was used to elongate each fragment to its original size using a custom pearl script, with the result stored as a BED file. Reads were then sorted and deduplicated such that only one fragment with the same starting and end position was retained. For visualization, wiggle files were generated with BEDTools version 2.23.0 (Quinlan and Hall, 2010), converted to bigwig format and analyzed in the Integrated Genome Browser (Freese et al., 2016). Peak detection was performed using MACS version 3.0.0b1 (https://github.com/macs3-project/MACS) using the elongated and deduplicated bed files as inputs and setting a q-value cutoff of 0.1.

### ATAC-seq

For ATAC-seq mapping, adapters were detected and removed using fastq-mcf of ea-utils (https://expressionanalysis.github.io/ea-utils, version 1.04.738) and mapped to chromosomes 1-19, X, Y and M of the mouse mm10 using bowtie (Langmead, et al., 2009) version 1.3.1), with the options ‘-y -m 1 -S -X 2000 --allow-contain’. Mapped paired-end reads were converted to a bed file by generating the original fragment using a custom perl script and duplicates were removed such that only one fragment with the same starting and end position was retained. Due to the repetitive nature of the Y chromosome and non-informative mitochondrial genome, reads mapped to either of them removed were and .wig files were generated using BEDtools and converted to bigwig format.

### Differential ATAC-seq Analysis

In order to identify regions with differential accessibilities between the NotoP and MP samples, we employed diffReps version 1.55.4 (Shen et al., 2013), using the elongated and deduplicated ATAC-NotoP and ATAC-MP samples as treatment and control respectively. We used the default parameters, with the exception of performing the statistical analysis using a G-test (‘--meth gt’) and disabling the DNA fragment shifting (“--frag 0”) due to using already elongated reads. This resulted in 16,680 regions that displayed either a significantly higher (more open in NotoP) or lower (more open in MP) accessibility. We then removed all those regions overlapping known promotors (mm10 UCSC refseq genes), defined as +/5kb from known transcription start sites, resulting in 13,890 differential regions.

### TF peaks overlaps

In order to avoid inclusion of TF peaks that fall within regions displaying mapping artefacts in ATAC-seq data, we removed all peaks falling within those 35 previously identified regions (Koch et al.,2017 and Buenrostro et al., 2015). We then overlapped the peak regions of the T-NMPdiff, T-NotoPdiff and Foxa2-NotoPdiff datasets to obtain all 7 possible combinatorial binding profiles. We then used the single bp maxima of each peak and intersected those with those ATAC regions displaying a significant increase in accessibility in NotoP cells. Finally, each peak was assigned potential target genes depending on the binding location. Intergenic peaks (at least 5kb away from any gene annotation) were assigned to both the closest up- and downstream gene, while genic peaks (those located between −5kb of the promoter and +5kb after the gene end) were assigned to that gene (or multiple in case of overlapping genes) as well as the closest up- and downstream gene.

### Heatmaps and Average Profiles

Heatmaps and average profiles were generated using SeqMINER (Ye et al., 2011). For plotting the ATAC-seq NotoP vs. MP comparison, the reference bed files were generated by using the center of differential ATAC regions and sorting them by the generated diffReps p-values. In the case of plotting the T, Foxa2, H3K27ac and ATAC profiles, we used the peak maxima of T-NMP^diff^, T-NotoP^diff^ or Foxa2-NotoP^diff^ (in that respective order if present) and sorted the peaks first by the 7 different combinations and then randomized the peaks within each category. The corresponding elongated and deduplicated bed files were used as inputs and hence set the extension size to 0.

### UpSet Plot

The UpSet plot was generated using UpSetR (Conway et al., 2017) to visualize associations between peak categories and associated genes.

## Data Availability

The accession number for the sequencing data reported in this paper is GEO: GSE235030

